# Cortico-muscular control of postural balance in older adults – effects of training on functional connectivity

**DOI:** 10.64898/2026.06.24.734264

**Authors:** Ruud A. J. Koster, Leila Alizadehsaravi, Jaap H. van Dieën, Sjoerd M. Bruijn, Nadia Dominici, Andreas Daffertshofer

## Abstract

**Background:** Balance training in older adults can lead to reduced centre of mass accelerations and reduced angular momenta after perturbations of unipedal stance, reflecting an enhanced ability to recover balance. It has been suggested that the co-occurring changes in muscle synergies indicated strategy-specific adaptations in feedback control.

**Methods:** We investigated the cortical involvement in such adaptations by focusing on the interaction between muscle synergies and cortical activity after perturbations. Twenty older adults (>65 years) underwent short-term and three-week long-term balance training, and we assessed their recovery from unpredictable mediolateral perturbations during unipedal stance. We measured high-density EEG and activation of leg and trunk muscles. The representations of the balance-related muscle synergies were localised in the cortex using coherence-based beamformers in the β-frequency band.

**Results:** Balance performance was accompanied by task-specific β-band activation in the somatotopic representation of the lower extremities in the primary motor cortex. The β-power significantly dropped during the response to perturbations, while the coherence with the activation of muscle synergies significantly increased, especially for synergies active in the early stage of balance recovery. The task-related changes in cortico-synergy coherence, especially during the later phase of balance recovery, were significantly affected by short-term training.

**Conclusion:** Refinements of feedback control seem to underlie balance improvements in older adults. The significant changes in the cortico-synergy interaction after balance training suggest cortical involvement in these refinements.

## 1. Introduction

The ability to maintain postural balance is paramount for many daily activities. The accompanying motor control requires a proper integration of multisensory information and a concurrent, coordinated activation of the musculature (Taube & Gollhofer, 2010; Torres-Oviedo & Ting, 2010). Both may be jeopardised with age, potentially resulting in the loss of independence, injuries, and even lethal accidents (Iosa et al., 2014; World Health Organization, 2007). How can older adults maintain postural balance and how can this be improved if needed? Many types of intervention that aim to improve balance have been investigated. Strikingly, where strength training fails (Boshuizen et al., 2005; Chandler et al., 1998), functional balance training may still improve balance in older adults (Lesinski et al., 2015). Several studies have claimed that functional balance training induces neural adaptations at several levels (Beck et al., 2007; Mynark & Koceja, 2002; Taubert et al., 2010). This suggests that improvements in balance may rely not only on muscular adaptations, but also on adaptations in motor control.

That balance improvement may capitalise on changes in motor control is not a new idea. Studies in young(er) adults revealed that balance training improves balance (Cadore et al., 2013; Gauffin et al., 1988; Halvarsson et al., 2013; Lesinski et al., 2015; Spiva et al., 2014). It induces both structural (anatomical) and functional adaptations in the central nervous system (Beck et al., 2007; Schubert et al., 2008; Taube et al., 2007; Taubert et al., 2010, 2016). Since these functional adaptations may manifest as reductions in reflex gain, they are thought to involve supraspinal, and possibly cortical mechanisms (Perez et al., 2004; Taube et al., 2007). Interestingly, Taube (2012) reported a training-induced reduction of cortical excitability that appeared to be associated with improvement in postural balance.

Similar experiments in older adults have failed thus far to show congruent mechanistic results (Alizadehsaravi et al., 2022; Ruffieux et al., 2017). While this questions whether balance ability in older adults can be improved by modulating reflexes, brain imaging indicates alternative forms of neural adaptations when balance improves. Balance training increases grey matter volume and changes white matter fractional anisotropy in a variety of brain regions (Taubert et al., 2010). Moreover, it increases the connectivity of (pre-)supplementary motor areas with both the prefrontal cortex and parietal regions, with the latter being associated with improvements in balance performance (Taubert et al., 2010, 2011). The increase in structural connectivity may also affect functional connectivity, thereby facilitating information transfer and yielding more suitable motor commands.

That adjustments in functional connectivity contribute to balance improvements gained recent support in a study on adaptations through feedback control in older adults (Koster et al., 2024). In this companion paper, we conjectured that muscle activation during postural balance may become more reliant on cortical motor commands after training because cortical adaptations allow for (fine-)tuning these commands to current task requirements. In the current study, we therefore expect to observe an increased cortical involvement after (successful) balance training. Such an increase in cortical involvement aligns well with the predictive capacity of cortical hemodynamics for balance performance, particularly in older adults (Lehmann et al., 2022).

Balance training may enact an upweighting of the cortical involvement via changes in functional cortico-muscular or cortico-spinal connectivity. Hence, in addition to changes in cortical activity itself, we expect balance training to increase functional cortico-muscular connectivity. Cortical motor commands are propagated along the pyramidal tract (Lemon, 2008), which can be characterised by a (partial) synchronisation of the sending and receiving neural populations (Brown, 2000; Murthy & Fetz, 1994; van Wijk et al., 2012b). This phenomenon can be quantified via cortico-muscular coherence (CMC), i.e. the degree of (phase) synchronisation between encephalographic and electromyographic signals. Several studies have demonstrated the relevance of changes in cortico-muscular coherence during balance tasks (Jacobs et al., 2015; Ozdemir et al., 2018; Varghese et al., 2019), including positive correlations between training-induced improvements in task performance and changes in cortico-muscular coherence (Perez et al., 2006).

The studies listed so far used single-muscle electromyography (EMG) as a proxy for the motoneuron activity (Conway et al., 1995). Postural balance, however, is thought to arise through modular control, by so-called muscle synergies (d’Avella et al., 2003), rather than controlling muscles individually (Torres-Oviedo & Ting, 2007, 2010). Balance control may therefore be better understood from a modular control perspective. Zandvoort et al. (2019) performed such an analysis and established, in young(er) adults, that selected functional synergy motor commands are manifested in the sensorimotor cortex during a balance task. To what extent this finding generalises to older adults is unclear, also because with increasing age the cortical drive to the muscles is weakened (Bayram et al., 2015).

In our companion paper (Koster et al., 2024) improvements in balance performance after training were accompanied by adaptations in kinematics and its underlying modular neuromuscular control. Here, we investigated whether these behavioural and neuromuscular adaptations were associated with changes in cortico-synergy coherence. We hypothesised that (1) postural balance in older adults involves cortico-synergy coherence, and (2) training-induced improvements in balance performance are accompanied by an increase in this cortico-synergy coherence.

## 2. Method

As said, we built on recent findings reported in our companion paper (Koster et al., 2024). In this earlier report we focused on training-induced changes in kinematics and muscle activity including muscle synergy composition and shape. Here we include the co-registered electro-encephalography (EEG), which allows for analysing the coherence between muscle synergy activation and cortical activity. However, first we briefly summarise the methods since they slightly deviate from the previous study.

### 2.1 Participants

Twenty older adults participated in this study (71.9 ± 4.1 years old; mean ± SD, 10 males, 9 favoured the left leg in unipedal stance). Participants were excluded if they presented with depression (GFS > 5), cognitively impairment (MMSE < 24), severe auditory or visual impairments, or if they were unable to stand and walk independently for at least 3 minutes. Additional exclusion criteria were obesity (BMI > 30); orthopaedic, neurological, and cardiovascular disease; and the use of balance affecting medication. Moreover, participants practicing sports that explicitly include a balance component were excluded to prevent ceiling effects (Kiers et al., 2013).

Participants were asked to maintain their normal activity levels in their daily life until the end of the experiment so as not to obscure intervention effects. The study was approved by the ethical review board (Faculty of Behaviour and Movement Sciences, Vrije Universiteit Amsterdam; VCWE-2018-171) and all participants provided written informed consent prior to participation.

### 2.2 Experimental procedure

The experiment consisted of a threefold measurement paradigm (repeated measures) with functional balance training in the interim (Fig. 1). Measurement session ‘Pre’ served as baseline, ‘Post1’ to assess short-term training effects, and ‘Post2’ to assess long-term training effects. A single measurement session consisted of registration of ‘resting state’ activity levels, task familiarization, followed by perturbed balance tasks. All balance trials took place on a custom-made robot-controlled balance platform (HapticMASTER, MOOG B.V., Nieuw-Vennep, The Netherlands). This platform could rotate 17.5° in both directions in the frontal plane. The robot arm could enact rotational perturbations or stimulate rotational stiffness.

**Figure 1:**
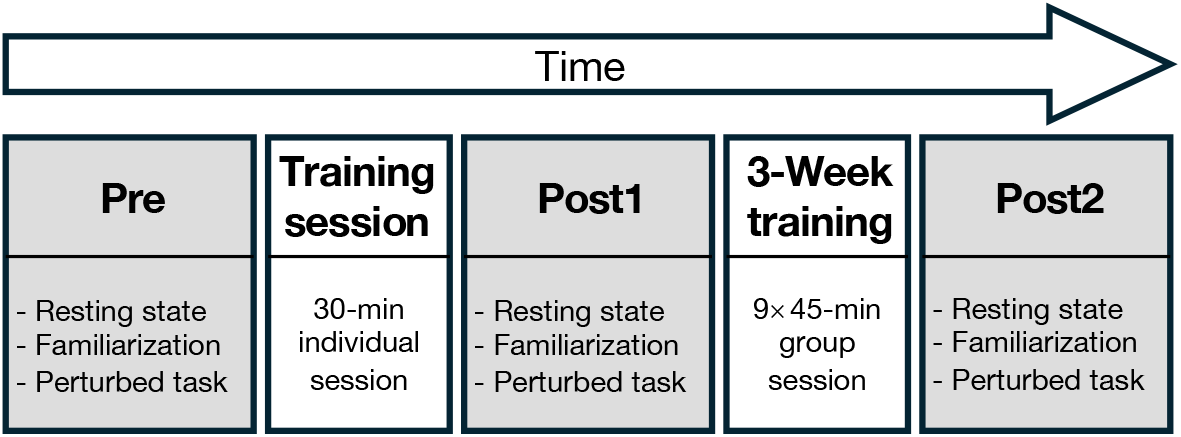
Overview of the protocol. Measurement session Pre was followed by 30 minutes of functional balance training and session Post1 on the same day. Session Post2 was preceded by a 3-week (9×45-min) balance training program. A session consisted of registration of resting state activity levels, task familiarization, followed by perturbed balance tasks.

‘Resting state’ activity levels were recorded while participants stood still on the floor in bipedal stance for 60 seconds. During balance tasks, participants were instructed to maintain unipedal stance on their preferred leg on the robot-controlled balance platform while keeping their arms abducted. First, participants underwent five familiarization trials during which the platform either had a constant stiffness or perturbed the participant slightly by rotating in the frontal plane (five times). Then, five trials of a perturbed balance task were performed. In each trial, twelve mechanical perturbations were imposed on the participant by rotating the platform. The perturbations (amplitude: 8°; max. angular speed: 16°/s) were smoothened parabolas, in a random initial direction and back, and with a random inter-perturbation interval of 3-5s to minimise anticipatory behaviour. Across analyses, we describe the perturbation direction relative to the standing leg: during lateral perturbations the inner side of the standing foot moved upward, tilting the participant lateral of the standing leg; during medial perturbations the inner side of the standing foot moved downwards. Inter-trial rest was granted to prevent fatigue.

During the first training session, participants were trained individually for 30 minutes. The three-week training program (9×45-min sessions) took place in groups of 6-8. The content of the training program was designed based on previous literature on balance improvement and fall risk reduction in older adults (Lesinski et al., 2015; Sherrington et al., 2017); see DOI:10.7717/peerj.18588/supp-1). Only standing balance exercises were included. All sessions were supervised by a physical therapist, who kept the exercises sufficiently challenging.

### 2.3 Data acquisition

Electro-encephalography (EEG) and electro-myography (EMG) were acquired during resting state and the perturbed balance task. EEG was recorded using a 64-electrode EEG cap (TMSi, Twente, The Netherlands) in accordance with the standard 10-20 montage. The impedance was kept below 20 kΩ by injecting impedance gel (SonoGel, Bad Camberg, Germany) between the electrodes and the scalp. The EEG signal was referenced to an average common reference, sampled at 2,048 Hz, and amplified using a 72-channel TMSi REFA amplifier (TMSi). Surface EMGs of 15 muscles were recorded using bipolar Ag/AgCl electrodes (∅ 11 mm, 20 mm inter-electrode distance, Ambu blue sensor N, Ambu, Ballerup, Denmark), placed in accordance with the SENIAM conventions (Hermens et al., 2000). Five bilateral hip and trunk muscles were recorded: rectus femoris (RF), adductor longus (AL), gluteus medius (GM), biceps femoris (BF), and erector spinae (ES) at the level of L2. Additionally, five muscles of the preferred stance leg were recorded: tibialis anterior (TA), soleus (SO), gastrocnemius lateralis (GL), peroneus longus (PL), and vastus lateralis (VL). The EMG signal was sampled at 2,000 Hz and amplified using a 16-channel TMSi PORTI system (TMSi).

### 2.4 Data analysis

#### 2.4.1 Pre-processing

All data processing and analysis was performed using Matlab (2021b; The MathWorks, Natick, MA, USA) and the FieldTrip toolbox for M/EEG analysis (Oostenveld et al., 2011). Throughout the analyses, filters were applied bi-directionally to preserve phase information.

*EMG*. The EMG data were up-sampled to 2,048 Hz to agree with the EEG, filtered (band-pass: 2^nd^ order Butterworth, 30-200 Hz; band-stop: 3^rd^ order Butterworth, 0.5 Hz bandwidth, *k*·50 Hz), and rectified using the absolute value of the Hilbert-transformed signal. From hereon we refer to this pre-processed EMG signal as the *broadband* EMG.

*EEG*. For the EEG, we defined signal outliers as samples that exceeded ±10 standard deviations (SD) from the channel’s mean and replaced them (as 2 ms time window) via linear interpolation. Bad EEG channels were repaired with spatially weighted integration of neighbours whenever possible. Channels were considered bad when their amplitude exceeded 10 times the mean absolute amplitude of all channels, or if their SD was 3× greater or 100× smaller than the mean SD of all channels (except for the channel to which the EEG was temporarily re-referenced). Thereafter, the EEG were re-referenced to a common average and filtered (band-pass: 2^nd^ order Butterworth, 1-200 Hz; band-stop: 3^rd^ order Butterworth, 0.1 Hz bandwidth, *k*·50 Hz). Subsequently, the EEG was decomposed using ICA. We removed independent components with a median frequency below 2 Hz (movement artefact) (Kline et al., 2015) and above 80 Hz (muscle activity) (Farella et al., 2002; Kroon et al., 1986; Kumar et al., 2001), and components that were dominated by pre-frontal channels (eye movements) (Jung et al., 1998) or by occipital and/or mastoid channels (neck muscle activity).

We normalised the EEG per participant and session to its mean total power during the 60s resting state recording obtained at the start of each recording session to improve reliability when averaging over participants and sessions. We also ‘standardised’ brain anatomy to the preferred standing leg. That is, EEG channels were left/right mirrored for participants who stood on their right leg. This approach implicitly assumes that the cortical processes underlying the task are symmetric between left- and right-leg stance conditions.

#### 2.4.2 Muscle synergy analysis

From the broadband EMG we estimated the signal envelopes that we used as a proxy for the net input to the motor unit pool (Boonstra & Breakspear, 2012; Bruns, 2004; Farina et al., 2004; Myers et al., 2003). For this, the broadband EMG was low pass filtered (2^nd^ order Butterworth, 5 Hz). The EMG envelopes were down sampled to 100 Hz to accelerate analysis. Per participant and session each EMG envelope was normalized to its mean. We extracted [-0.5, 2.5]s epochs around perturbation onset and computed the average EMG envelope per participant, session, and perturbation direction. These EMG envelopes were expressed as perturbation-related activity by subtracting the mean activity in the last 0.5s before perturbation onset for the corresponding session and perturbation direction.

After concatenating the average EMG envelopes from all participants, sessions, and perturbation directions, principal component analysis (PCA) served to estimate muscle synergies, consisting of muscle weighting factors and a temporal activation pattern^1^. The number of synergies was selected such that at least 70% of the total variance in the concatenated EMG envelopes was explained, resulting in the extraction of 5 synergies. The resulting muscle weighting factors indicate the spatial structure of the modular control, i.e., how much a synergy contributes to the activation pattern of a particular muscle, whereas the temporal activation patterns represent each synergy’s general activation intensity over time. To also obtain the broadband temporal synergy activation patterns for each individual perturbation, i.e., the signal representing the modular motor command, the broadband EMG was multiplied by the inverse of the muscle weighting factors. Finally, the synergy time series, along with the EEG time series, were partitioned into [-0.5, 2.5]s epochs around perturbation onset as before.

The outcome of the EMG synergy analysis has been extensively discussed in the companion paper. Hence, we here refer to (Koster et al., 2024) for more details and corresponding interpretations. As reported in the companion paper, training effects were most pronounced in synergy 1, which explained the largest proportion of signal variance. As such, we will focus on this primary synergy in this paper and refer to the *Appendix* for the other synergies. We depict the corresponding activation patterns and muscle weightings in Figure 2 (the statistics is briefly summarised in *Appendix* A1).

**Figure 2:**
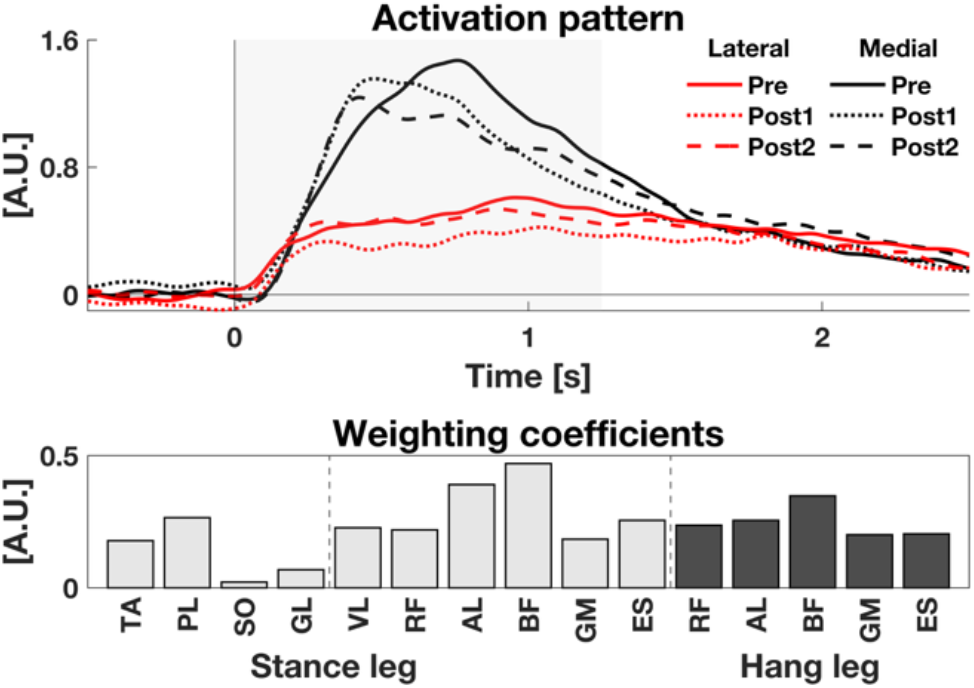
Synergy 1. *Top panel*: Mean temporal activation patterns for lateral (red) and medial (black) perturbations across sessions (Pre = solid lines, Post1 = dotted lines, Post2 = dashed lines). Baseline activity has been normalised to zero. *Bottom panel*: Corresponding muscle weighting factors obtained using PCA after concatenation of EMG envelopes across participants, sessions, and perturbation directions. Muscle abbreviations: RF = rectus femoris, AL = adductor longus, GM = gluteus medius, BF = biceps femoris, ES = erector spinae, TA = tibialis anterior, SO = soleus, GL = gastrocnemius lateralis, PL = peroneus longus, VL = vastus lateralis. Statistical results are summarised in *Appendix A1*.

##### Epochs-of-interest

As shown *Appendix* A1, the activation pattern of synergy 1 showed significant training effects during the early, dynamic, response phase and the late, static, phase. In what follows, we will use the spectral activities in these dynamic early, [0.1, 0.4]s, and static late response phases, [0.7, 1.0]s, and contrast them to the equally long pre-perturbation epoch [-0.4, -0.1]s.

#### 2.4.3 Spectral analysis

To compute the cortico-synergy coherence time-frequency analysis was performed on the filtered EEG data and broadband synergy activation patterns. Using Morlet wavelets (relative bandwidth = 3), we determined the following cross-spectrum matrices: *C*_EEG,EEG_ (between all EEG electrodes), *C*_EEG,ref_ (between the EEG electrodes and the reference synergy 1 activation patterns), and *C*_ref,ref_ (synergy 1 with itself). We used the aforementioned epochs-of-interest and focused our analysis on the β-frequency band, [15, 30] Hz: a frequency range in which cortico-muscular coherence (Kristeva et al., 2007; Reyes et al., 2017; Roeder et al., 2018), and more recently cortico-synergy coherence (Koster et al., 2025; Zandvoort et al., 2019, 2022), has been observed.

The β-band is commonly associated with motor control (Engel & Fries, 2010; Gross et al., 2000; Mima et al., 2000). However, since EEG during perturbed balance tasks in older adults has been only sparsely investigated, we first performed an exploratory analysis to confirm the relevance of the β-band for the present task. Figure 3 shows the spectral power as a function of time and frequency averaged of 9 central/mesial EEG channels that arguably record the lower extremity representations in bilateral primary motor cortices. For the β-band one can clearly observe a drop in the spectral power above the cortical representation of the lower extremities after perturbation onset at *t* = 0s. Such a drop in cortical β-power, also called event-related β-desynchronisation, is characteristic of dynamic motor responses (Neuper & Pfurtscheller, 2001; Seeber et al., 2014; van Wijk et al., 2012a). This pre-analysis corroborated our focus on the β-frequency band and, further supported our choice for the epochs-of-interest as [-0.4, - 0.1]s for the baseline and [0.1, 0.4]s and [0.7, 1.0]s for the dynamic early and static late responses, respectively. Note that in line with Figure 2, left top panel we always normalised the baseline log β-power by subtracting the mean value over the [-0.4, -0.1]s by which the baseline was not only steady over time but differences between subjects were also minimised.

**Figure 3:**
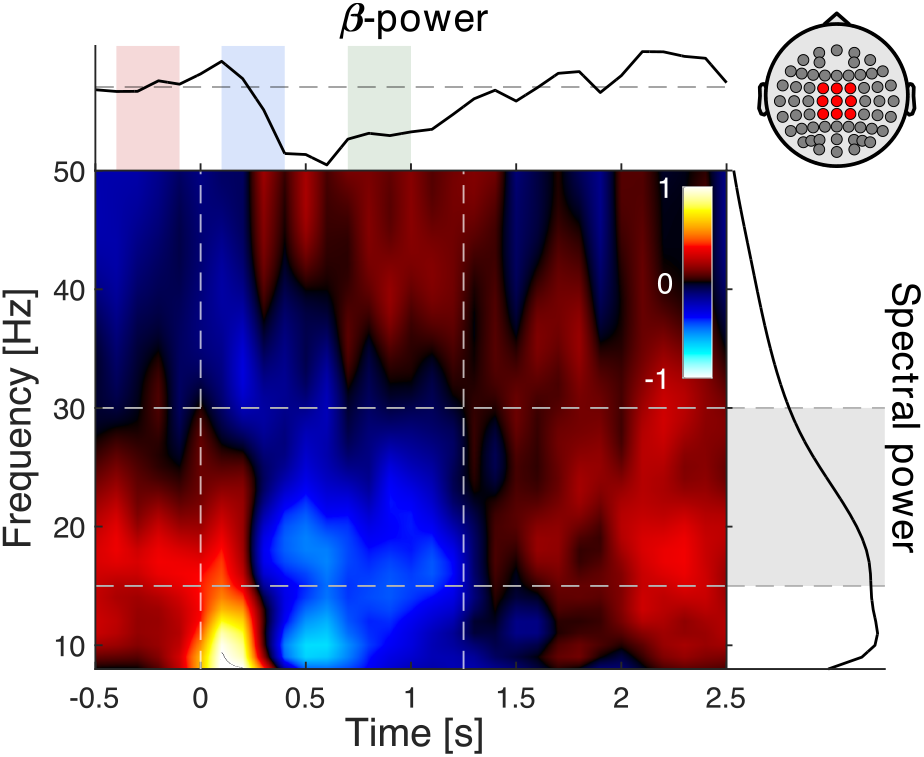
Time-frequency representation of central EEG power, normalised to the baseline, pooled over participants, session, and perturbation direction. After perturbation onset at 0s an induced response is clearly visible in the β-frequency band [15-30] Hz. This response is marked by a substantial drop in spectral power characteristic for dynamic motor activity and has a duration comparable with the kinematic responses. The coloured areas in the top plot indicate the baseline and early and late response epochs. The grey area in the right plot indicates the β-band. The vertical lines in the time-frequency plot indicate the epoch of interest in which the kinematic response occurred.

#### 2.4.4 Source analysis

To assess cortico-synergy coherence, we identified the cortical sources associated with the balance task and estimated their (virtual) activation. We briefly summarise our approach to this.

##### Source localisation

We first constructed a forward model, which maps activity from any location in the cortex to the EEG sensors. We used a five-layer volume conduction model including white and grey matter, cerebrospinal fluid, skull tissue, and scalp tissue. Anatomy was based on the “Neurodevelopmental MRI Database” MRI template for 70-74 years old adults (Richards et al., 2016). We used finite element modelling to restructure the volume conduction model into a realistically shaped volumetric grid using the following conductivity values (in S/m): white matter = 0.14, grey matter = 0.33, cerebrospinal fluid = 1.79, skull = 0.0, and scalp = 0.43 (Birot et al., 2014). This allowed for constructing a lead field matrix and subsequently a source model at a 2 mm^3^ voxel resolution.

We estimated spatial filters via dynamic imaging of coherent sources (DICS) beamformers (Gross et al., 2001), which incorporates the cross-spectra between the EEG channels. This allowed for projecting the EEG sensor activity to each individual voxels, i.e. to every (virtual) source *ν*. For the resulting source activities, we estimated the (squared) coherence with our reference activity of synergy 1 using:

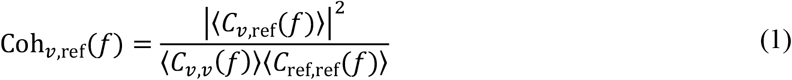

where *C*_*ν*,ref_(*f*) denotes the cross-spectrum between the virtual activity at voxel *ν* and the activity of synergy 1, and *C*_*ν*,v_(*f*) and *C*_ref,ref_(*f*) represent the respective auto- or power-spectra; ⟨·⟩ indicates the expectation value that we determined by averaging over all perturbations irrespective of their direction; and *f* is the frequency of interest (here the β-band).

The coherence was determined for every voxel in the cortex for every participant and session; the subsequent statistical evaluation can be found below under *Statistics*.

##### Source reconstruction

The statistical evaluation of the voxel-wise coherence yielded task-related cortical sources. For these voxels we set spatial filters by averaging corresponding cross-spectra over sessions. In contrast to the definition above we here optimised the filters for the epoch [0, 1.25]s after perturbation onset, to focus on the full epoch where the kinematic response took place, also to be consistent with the analyses in our companion paper (Koster et al., 2024). Per participant, a single source-level spatial filter was defined by weighted averaging all its voxel-level spatial filters. As weighting factor, we used the absolute value of a voxel’s *t*-statistic for the baseline-activity contrast, i.e. the more the activity of a voxel was task-related, the more that voxel contributed to the overall source activity (see *Statistics* below). For the resulting source activities, we estimated the spectral power using the same approach we used at the sensor-level analysis. For each participant, session, and perturbation direction we also estimated the β-band cortico-synergy coherence over time using Eq. (1).

### 2.5 Statistics

#### Source localisation

To identify cortical regions that were involved in the balance task, we used two contrasts: the early dynamic response epoch versus the baseline epoch, i.e. [0.1, 0.4]s vs [-0.4, -0.1]s, and the late static response epoch versus the baseline epoch, i.e. [0.7, 1.0]s vs [-0.4, -0.1]s. To warrant consistency of the respective statistical estimates we always used a ‘common’ spatial filter that spanned both epochs. Note that we assumed source location invariant over recording sessions and used each epoch’s average coherence over sessions per participant (see also below for the corresponding verification).

Per contrast, we conducted paired *t*-tests (baseline vs activity, over participants) for every voxel (*n* = 62437). To mitigate family-wise error rate inflation, cluster-based Monte Carlo permutation testing was performed (Ernst, 2004; Maris, 2012; Maris et al., 2007; Maris & Oostenveld, 2007; Pernet et al., 2015). Clusters were defined as adjoining voxels with *t*-test’s *p*-values below *α*_*t*-test_= 0.05. For each cluster, we defined its statistics *M*_true_ as the sum of its voxels’ *t*-statistics. This computation was repeated for 10,000 random permutations (for every voxel and participant the ‘baseline-activity’-value pair was randomly switched places). For each permutation *i* we determined the maximum cluster-statistic *M*_*i*_ and constructed the null distribution using {*M*_1_, …, *M*_*i*_, …, *M*_10,000_}. The statistical test’s *p*-value was finally defined as the proportion of *M*_*i*_ whose value is more extreme than *M*_true_ of each cluster in the true data and evaluated using the significance threshold *α*_cluster_ = 0.05.

Only sources in the bilateral pre- and postcentral gyri of the LONI Probabilistic Brain Atlas (LPBA40) (Richards et al., 2016; Shattuck et al., 2008) were considered in the next step of the analysis. The cortical-wide statistical evaluation served as a useful check of common assumptions: that cortico-muscular (or cortico-synergy) coherence was primarily localised within sensorimotor regions. Nevertheless, restricting the analysis to the pre-defined region of interest was necessary to ensure that the identified voxels represented a functionally coherent source. This was required for subsequent averaging of activity over the entire source.

Whenever cortical sources were identified, the cluster-based permutation test was also performed based on a one-way repeated-measures ANOVA (‘sessions’ as factor with three levels: Pre, Post1, and Post2), for which coherence was not averaged over sessions. This primarily served to verify the assumption that the source location was invariant over recording session, i.e., validating the reliability of the source.

#### Source activity

To evaluate task-related cortical activity we contrasted the log β-power and cortico-synergy coherence during baseline against early and late responses epochs. For this we used two-tailed paired *t*-tests and pooled the data over perturbation directions and recording sessions. For the effect of training, we determined the corresponding mean differences of the power and the coherence (early response-baseline and late-response-baseline epoch) and conducted one-way repeated-measures ANOVAs with three levels: Pre, Post1, and Post2. For significant differences between sessions, we conducted post-hoc two-tailed paired *t*-tests (Pre-Post1, Pre-Post2, Post1-Post2) with Bonferroni correction. In line with the synergy analysis in our companion paper (Koster et al., 2024) we always normalised to the baseline log β-power and cortico-synergy coherence by subtracting the mean value over the [-0.4, -0.1]s per subject prior to hypothesis testing.

We also compared the β-power and cortico-synergy coherence time series, for medial and lateral perturbations separately. The epoch over which these statistical analyses were conducted was [0, 1.25]s after perturbation onset. Again, we used a one-way repeated-measures ANOVA with three levels (the sessions), but given the dimensionality (time-resolved) we here employed statistical parametric mapping (SPM) (Pataky et al., 2016). If significant differences between sessions were present, the ANOVAs were supplemented by post-hoc two-tailed paired *t*-tests with Bonferroni correction.

Throughout hypothesis testing the significance threshold was set to *α* = 0.05.

## 3. Results

### 3.1 Source localization

During the balance task, coherence between the source-reconstructed EEG and the activation pattern of synergy 1 was observed central bilaterally in the motor cortex, extending slightly into pre-motor and supplementary motor areas. As illustrated in Figure 4, the cortico-synergy coherence was elevated in the early response phase, relative to the pre-perturbation baseline. This increase reached statistical significance, but only in the left hemisphere, and only when the data of all three sessions were pooled (*t*_max_ = 3.52; *p* = .0034). Separating the data of the three session and conducting an ANOVA did not reveal significant differences between the Pre, Post1, and Post2 sessions. These findings suggest that the localisation of the task-related source remained stable across training sessions, supporting the validity of pooling the data. Evaluating source-level coherence for medial & lateral perturbation directions individually did not yield significant sources, potentially due to the reduced sample size (i.e., 30 perturbations per participant/session/direction) – we return to this in the *Discussion* section.

**Figure 4:**
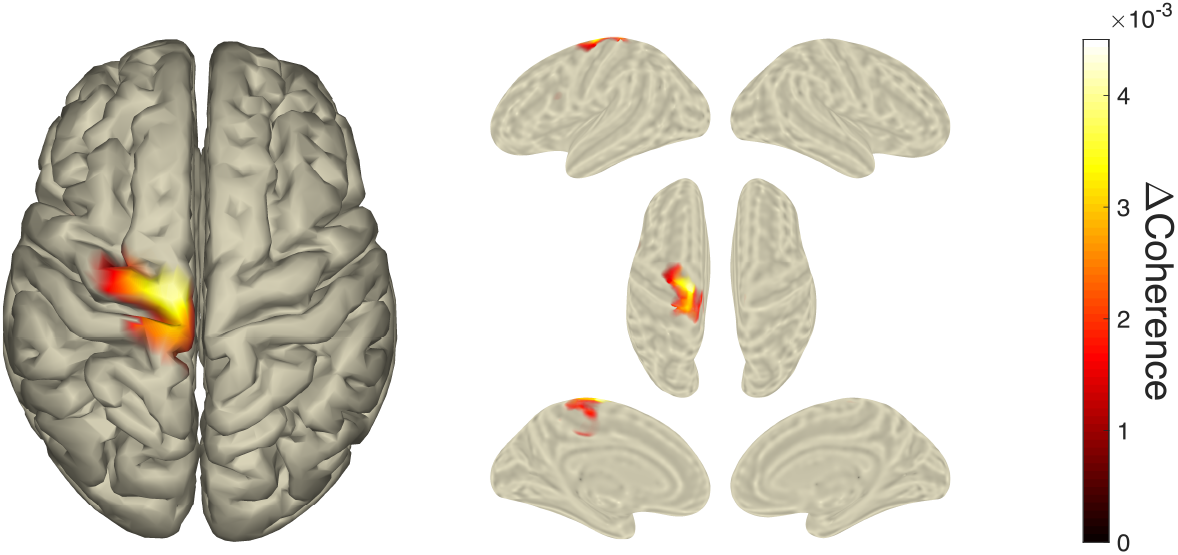
*Left panel*: Top-view cortical surface showing changes in cortico-synergy coherence between the baseline and early response epochs, pooled over sessions. *Right panel*: Corresponding inflated cortical surface representation. Coherence maps are masked for statistical significance; the unmasked maps are shown *Appendix A2*.

During the late response phase, the maximum cortico-synergy coherence presented around the (bilateral) motor cortex (see *Appendix* A2). In contrast to the early response, however, the coherence did not differ significantly from pre-perturbation baseline.

For an overview of the cortico-synergy coherence of the other synergies, we refer to *Appendix A3*. Here we only add that, although some voxels reached significance, none formed spatially coherent sources within the expected motor network, and that the coherence levels of the other synergies were only a fraction of those observed in synergy 1.

### 3.2 Source activity

The activity of the identified cortical source revealed temporal changes in spectral power that largely agreed with the sensor-level analysis; cf. Figure 3 & Figure 5 (left panel). Here, only the significant voxels (within the pre- or postcentral gyri) were considered, weighted according to their task-relevance for optimal source reconstruction. The time-resolved cortico-synergy coherence also displayed an event-related response, with an increase in β-band coherence shortly after perturbation onset that lasted until approximately 1.25s after perturbation onset. In line with Figure 4, the coherence increased most in the early response epoch [0.1, 0.4]s.

**Figure 5:**
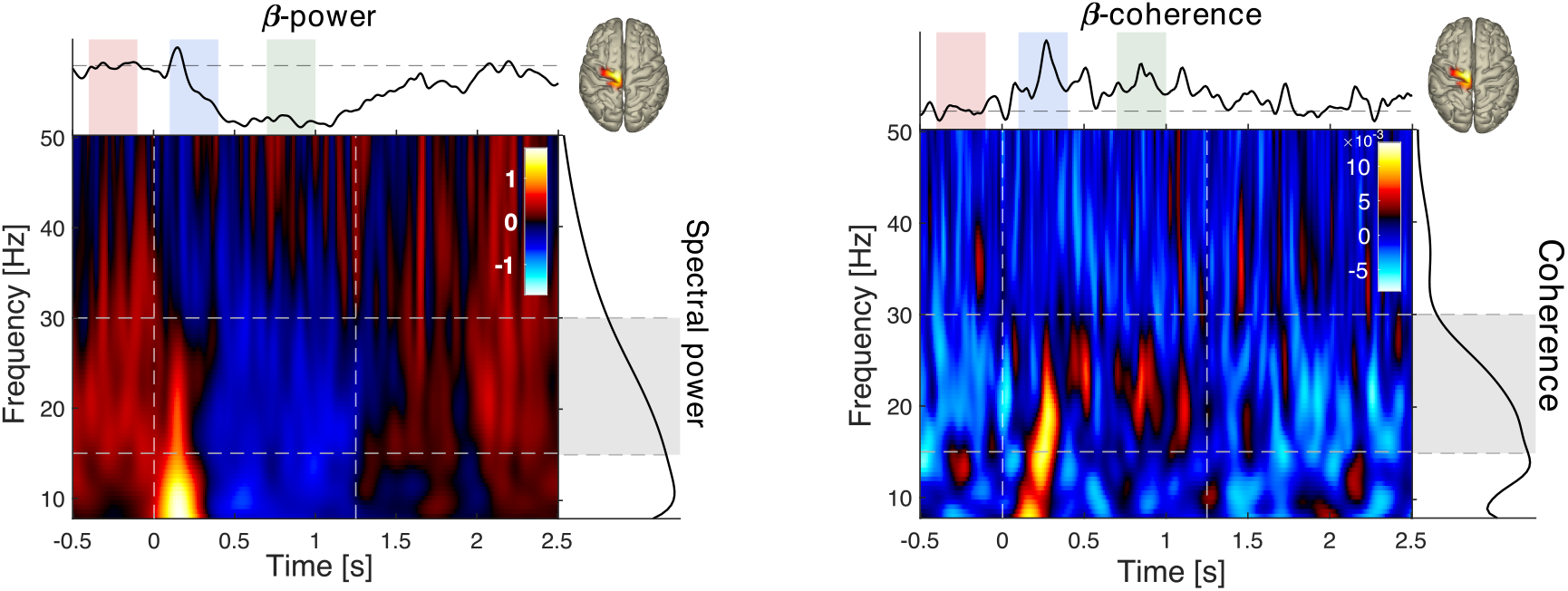
Time-frequency representation of source level power (*left panel*) and cortical-synergy coherence (*right panel*). Data have been pooled over session and perturbation direction. The top graphs show the respective mean values over the β-frequency band. For the spectral power, this reveals the characteristic event-related desynchronization shortly after perturbation onset, coinciding with an increase in synergy/muscle activation. The graphs right to the time-frequency plots show the log-power (*left panel*) and coherence (*right panel*) averaged over time; cf. Figure 3 for further details.

These temporal changes in power and coherence supported our contrast definitions within the [15-30] Hz frequency band. Specifically, the interval[-0.4, -0.1]s represented a stable baseline epoch, [0.1, 0.4]s captured the highly dynamic early response, and [0.7, 1.0]s corresponded to the stable late response. Moreover, both the event-related drop in power and the accompanied rise in coherence align remarkably well with the [0, 1.25]s kinematic response epoch reported in Koster et al. (2024).

The descriptive statistics of the spectral distributions pooled across sessions and perturbation directions suggested an event-related drop in β-power together with an increase in coherence that lasted approximately 1.25s after perturbation onset (Figure 5). This was confirmed by the *t*-tests that revealed reductions in β-power and increases in coherence. The power changes reached significance during the late response epoch, while coherence changed significantly during both response epochs (see Table 1).

**Table 1.** Comparison between the mean baseline and response activity after pooling the data over perturbation directions and sessions (*df* = 19).

|  | baseline vs early response |  | baseline vs late response |  | late response vs early response |  |
| --- | --- | --- | --- | --- | --- | --- |
| | $t$-value | $p$-value | $t$-value | $p$-value | $t$-value | $p$-value |
| power | -1.34 | .196 | -4.80 | **<.001** | -3.86 | **.001** |
| coherence | 3.83 | **.001** | 3.22 | **.005** | -0.36 | .721 |

As summarised in Table 2, the task-related changes in coherence during the late response were significantly affected by training. The post hoc *t*-tests indicated that the difference in coherence between baseline and late response was highest in Post1 (Pre-Post1: *t* = 2.41, *p* = .026; Pre-Post2: *t* = -0.56, *p* = .581; Post1-Post2: *t* = -3.77, *p* = .001).

**Table 2.** Comparison between sessions of the mean difference between baseline and response activities pooled over perturbation directions (*df* = 2,38).

|  | baseline vs early response |  | baseline vs late response |  | late response vs early response |  |
| --- | --- | --- | --- | --- | --- | --- |
|  | *F*-value | *p*-value | *F*-value | *p*-value | *F*-value | *p*-value |
| power | 0.67 | .516 | 0.16 | .838 | 0.23 | .747 |
| coherence | 0.91 | .412 | 6.19 | **.006** | 0.87 | .425 |

Without averaging over sessions, however, the evidence for an event-related rise in coherence was less clear and coherence values showed considerable variability time (Figure 6). Yet, the comparison of the time-resolved cortico-synergy coherence between sessions indicated a significant main effect of training, both for the mean β-power and for the cortico-synergy coherence (Figure 6). For the latter, post-hoc testing revealed significant differences between the Post1 and Post2 sessions during the late response. However, when assessed separately for each perturbation direction, these effects did not reach statistical significance (*Appendices A1* and *A4*).

**Figure 6:**
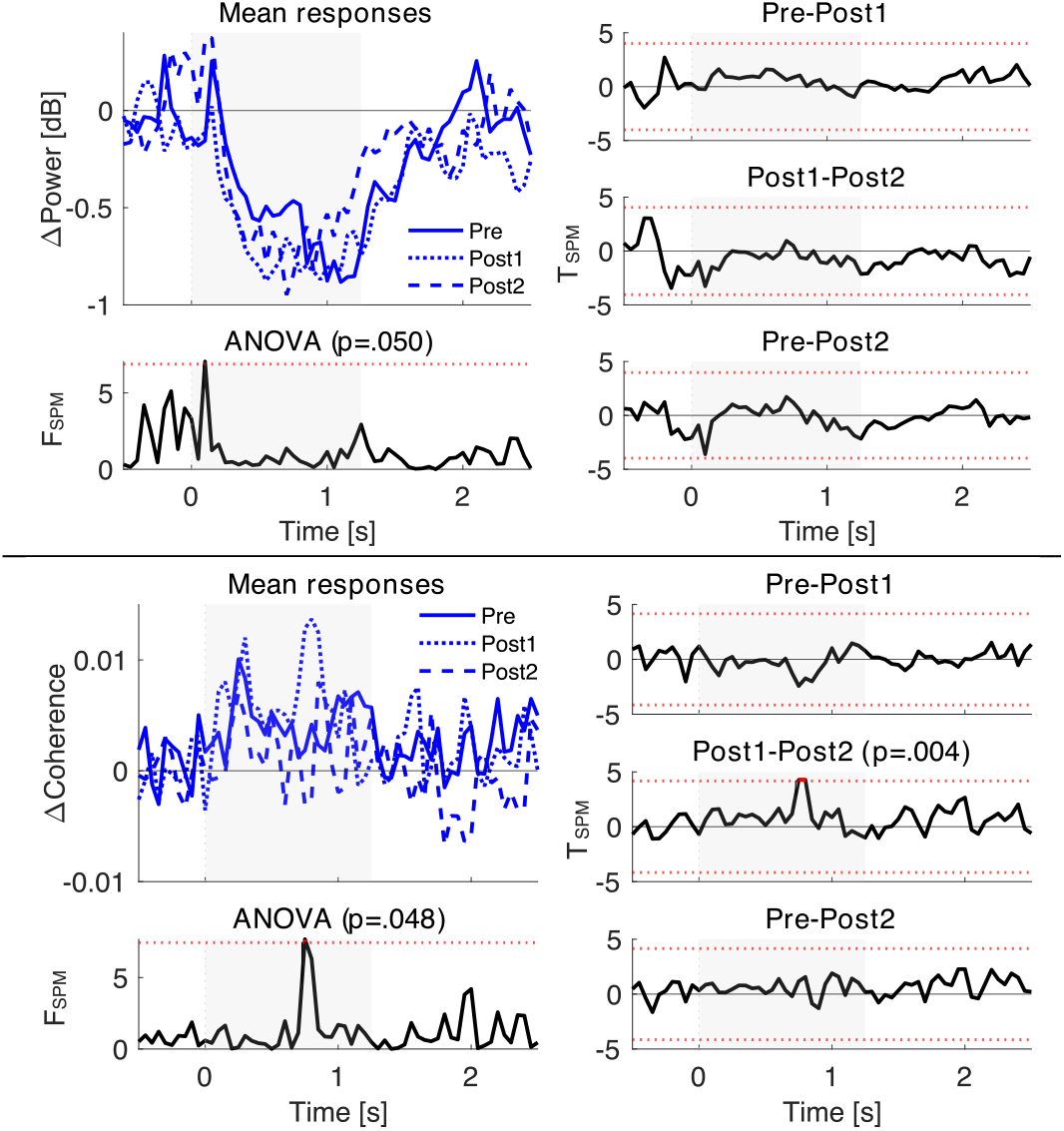
Between sessions comparison of the β-power (*top panel*) and the cortico-synergy coherence (*bottom panels*) pooled over perturbation directions. Top rows (per panel): mean power/coherence as a function of time (*t* = 0s ∼ perturbation onset). Bottom rows (per panel): the corresponding *F*-statistics, the red dotted line indicates the significance threshold. For each participant, baseline log-power or coherence values were subtracted to reduce between-subject variability. Therefore, power and coherence are represented as ΔPower and ΔCoherence, respectively.

See *Appendix A3* for the results per perturbation direction.

## 4. Discussion

In the current study, we built on our companion paper in which we established that balance training can improve balance performance in older adults. This included adaptations in the kinematics and the underlying modular neuromuscular control (Koster et al., 2024). Here we expected that postural balance in older adults would involve cortico-synergy coherence, as had previously been observed in younger adults (Zandvoort et al., 2019). Our findings confirmed this both at sensor level and, even more prominently, at the cortical source level. Figure 4 clearly displays a statistically significant, focal coherence in the β-frequency band in the primary motor cortex representation of the non-standing leg.

From a utility perspective this lateralisation towards the non-standing leg is not a complete surprise. van Dieën et al. (2015) demonstrated that, in unipedal stance, the non-stance leg is the largest contributor to the corrective change in angular momentum, making its control crucial for postural balance.

Cortical changes in the β-band, especially motor-related desynchronisation, are seminal for dynamic motor activity (Kilner et al., 1999; van Wijk et al., 2012b). As such, the non-stance leg appeared to compensate both the medial and lateral perturbations, and the observed changes in spectral power and coherence are indicative of its underlying motor processing. Nonetheless, since synergy 1 governs muscles of both legs it is striking that the identified source location did not extend into the motor cortex representations of the standing leg in the other hemisphere. The descriptive statistics of coherence (see *Appendix A2)* show little evidence of lateralisation to either hemisphere. Given our cluster-based statistical approach, where a cluster is either included or excluded in its totality, there remains a possibility that the observed lateralisation has a statistical rather than functional basis.

The task-related changes in cortico-synergy coherence in the β-band were significant both for the early and the late response epoch, while the difference in β-power reached significance only in the late response epoch (Table 1). A drop in power potentially reduces the signal-to-noise ratio that may decrease but not increase a coherence estimate. This suggests that the task-related changes in coherence were not just epiphenomenal to the changes in spectral power, which, as expected, dropped during dynamic responses relative to the steady baseline activity.

Training affected balance-related coherence. The ‘late response vs. baseline’ contrast differed significantly between sessions when averaging coherence over the epochs (Table 2). A closer look at the time-resolved changes confirmed this, indicating coherence was increased most after short-term training (Figure 6). The earlier reported training effects on the synergy’s activity (Koster et al., 2024), especially in the later balance response, seem to be matched by training-induced changes in cortico-synergy coherence. Yet, where synergy activity was affected by both short- and long-term training, its cortico-synergy coherence appeared to only be affected by short-term training. Moreover, when looking at the perturbation directions separately, as done in Koster et al., (2024), the changes in coherence did not pass our significance threshold. Several factors may explain these findings, with the limited sample size being an important consideration.

Cortico-muscular coherence or cortico-synergy coherence values were expected to be very small during balance performance, especially after a perturbation. In our analyses, the coherence values hardly exceeded 0.1 (note that we reported the changes with respect to baseline which are even smaller). That is, the coherence was observed in a range which is known to be particularly prone to an estimation bias when the sample size is small. As mentioned in the *Results* section 3.1, we used 30 perturbations per participant/session/direction which may appear to be a lot in the experimental design, but it implies that coherence was strongly positively biased, i.e. the true coherence values were likely smaller than 0.1. Since this bias was constant across conditions, our statistical evaluation did not contain a systematic error. Nonetheless, any potential differences would be underestimated because of the bias flattening the coherence estimate (Amos & Koopmans, 1963; Carter & Nuttall, 1972; Enochson & Goodman, 1965). Combining this with a low signal-to-noise ratio and a substantial within- and between-subject variability, possible effects of balance training on coherence may therefore have remained undetected.

A similar note can be placed regarding our principal EEG analysis and its statistical evaluation. While the EEG source-localisation we used is standard, we addressed the concomitant multiple comparison problem through family-wise error correction to guarantee statistical rigour. By this choice, the source that we identified was stable over experimental sessions, albeit within the predefined region-of-interest. However, cortical contributions from other regions or frequency bands may have remained undetected, including alternative mechanisms underlying EEG-EMG coherence (Mima & Hallett, 1999).

As said, we did find significant effects of training on the cortico-synergy coherence. The effect size, however, was small and we rather abstain from interpreting the precise moment of change in more detail. Yet, the effect was largest in the Post1 session, i.e. after short-term training, which suggests that training duration and intensity might have been suboptimal to cause long-lasting changes in cortico-muscular/cortico-spinal connectivity or upweighting of the cortical input in the formation of motor output.

Despite the limitations our results do support the idea that training-induced balance improvement including altered kinematics and synergy activity is facilitated by cortical adaptations that may support the generation of more effective motor commands. Cortico-muscular coherence is a common means to quantify the functional interaction between cortex and spinal cord (and/or muscle activity). As shown here as well as in many other circumstances, it modulates with the task under study, which strongly suggests that this coherence quantifies specific type of functional coupling in the motor system (Liu et al., 2019; Mima & Hallett, 1999). However, the extent to which these training-induced balance improvements in older adults are mediated by cortical adaptations remains to be determined.

## 5. Conclusion

One-legged balance performance in older adults was associated with focal cortico-synergy coherence within the β-frequency band in the somatotopic leg representation of the primary motor cortex. These findings are consistent with previous observations in younger adults and align with kinematic responses involved in compensating for postural perturbations. Moreover, balance training altered cortico-synergy coherence alongside improvements in balance performance, supporting the idea that cortical adaptations contribute to training-related refinements in postural control.

## Acknowledgement

This project received funding from the European Union’s Horizon 2020 research and innovation programme under the Marie Skłodowska-Curie grant agreement No 721577. RAJK was supported by the European Research Council (ERC) under the European Union’s Horizon 2020 Research and Innovation Program (grant agreement number 715945 Learn2Walk).

## Appendix

### A1 Statistics of muscle synergy 1

Figure A1 summarises the SPM-based ANOVA and the corresponding post-hoc assessments reported earlier in the companion paper (Koster et al., 2024) and (Koster et al., 2026).

**Figure A1.**
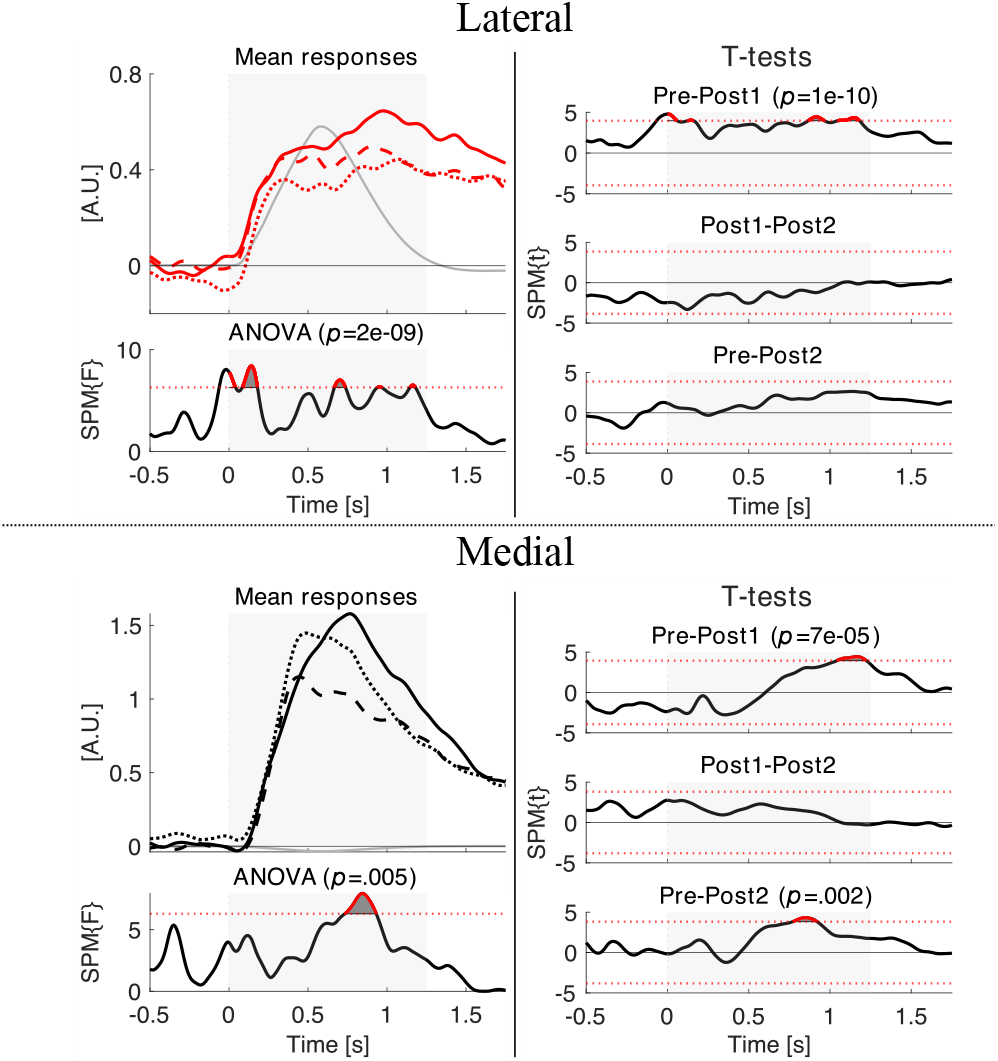
Statistics of muscle synergy 1. *Top panel*: lateral perturbation; *bottom panel*: medial perturbation. *Left column*: (*top*) mean temporal activation patterns per session (mean of the average per participant; Pre = solid lines, Post1 = dotted lines, Post2 = dashed lines), (*bottom*) *F*-statistics of the corresponding ANOVA comparison between sessions. The horizontal red dotted line indicates the significance threshold. *Right column*: *t*-statistic of post-hoc comparisons between sessions. The horizontal red dotted lines indicate the significance threshold. The grey curve in the left top plot indicates the shape and timing of the perturbation expressed as platform rotation. The ANOVA results (*F*-statistics) served to define the epochs of interest as [0.1, 0.4]s (early response) and [0.7, 1.0]s (late response) that were contrasted to the pre-perturbation baseline [-0.4, -0.1]s.

### A2 Cortico-synergy coherence – early and late responses

**Figure A2.1.**
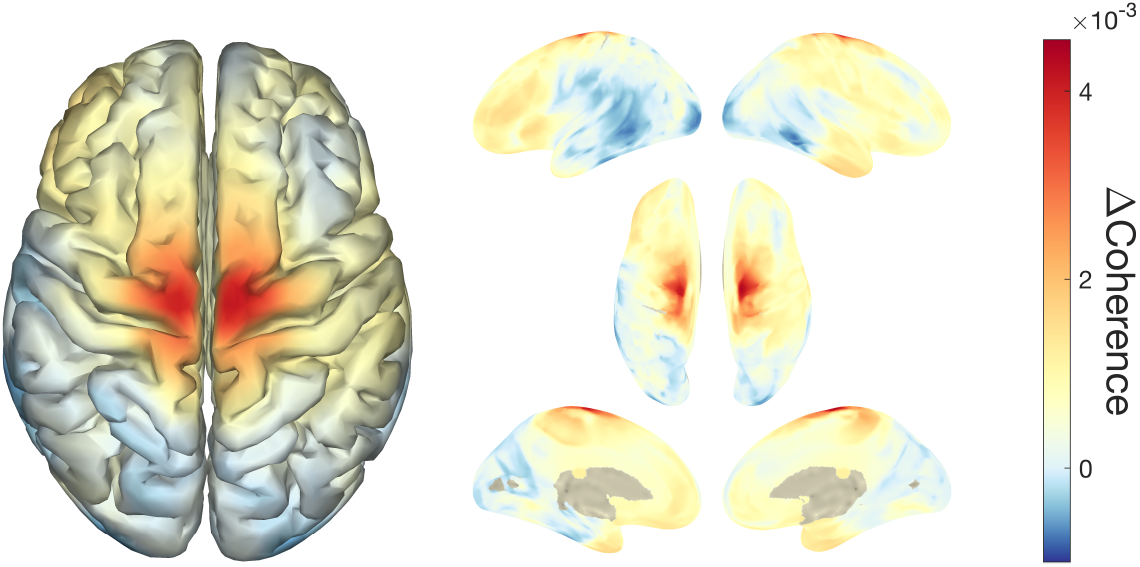
Left panel: Top-view surface plot of the differences between the cortico-synergy coherence before-after perturbation (early responses, pooled over sessions). Right panel: idem but using an inflated cortical surface.

**Figure A2.2.**
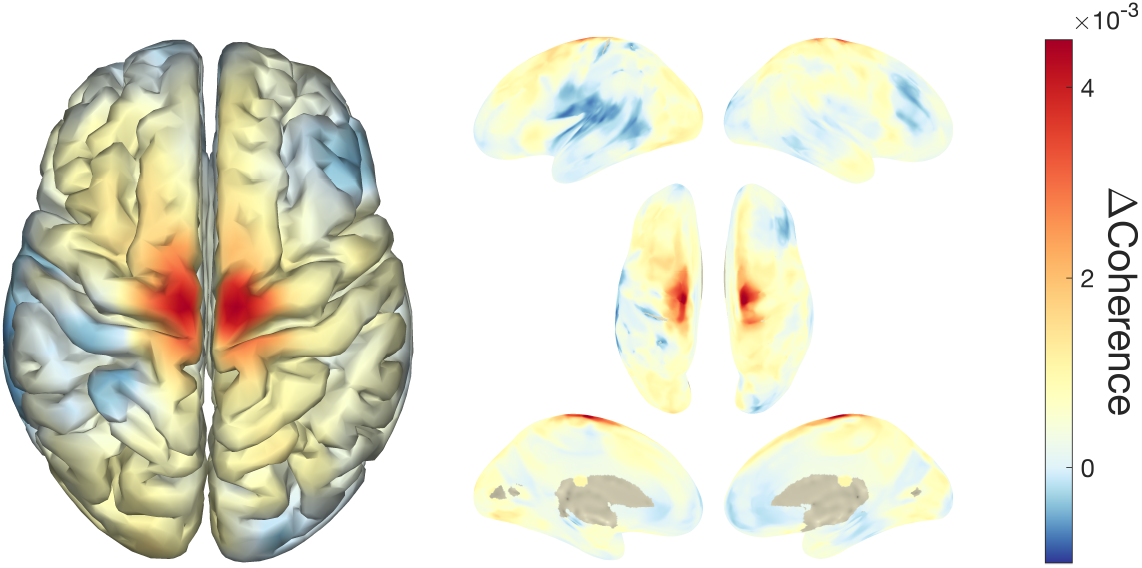
Same as Fig. A2.1 but for the late responses, pooled over sessions). Right panel: idem but using an inflated cortical surface. Note that no statistically significant sources could be identified in this contrast.

### A3 Cortico-synergy coherence estimates of all synergies

**Figure SM1.**
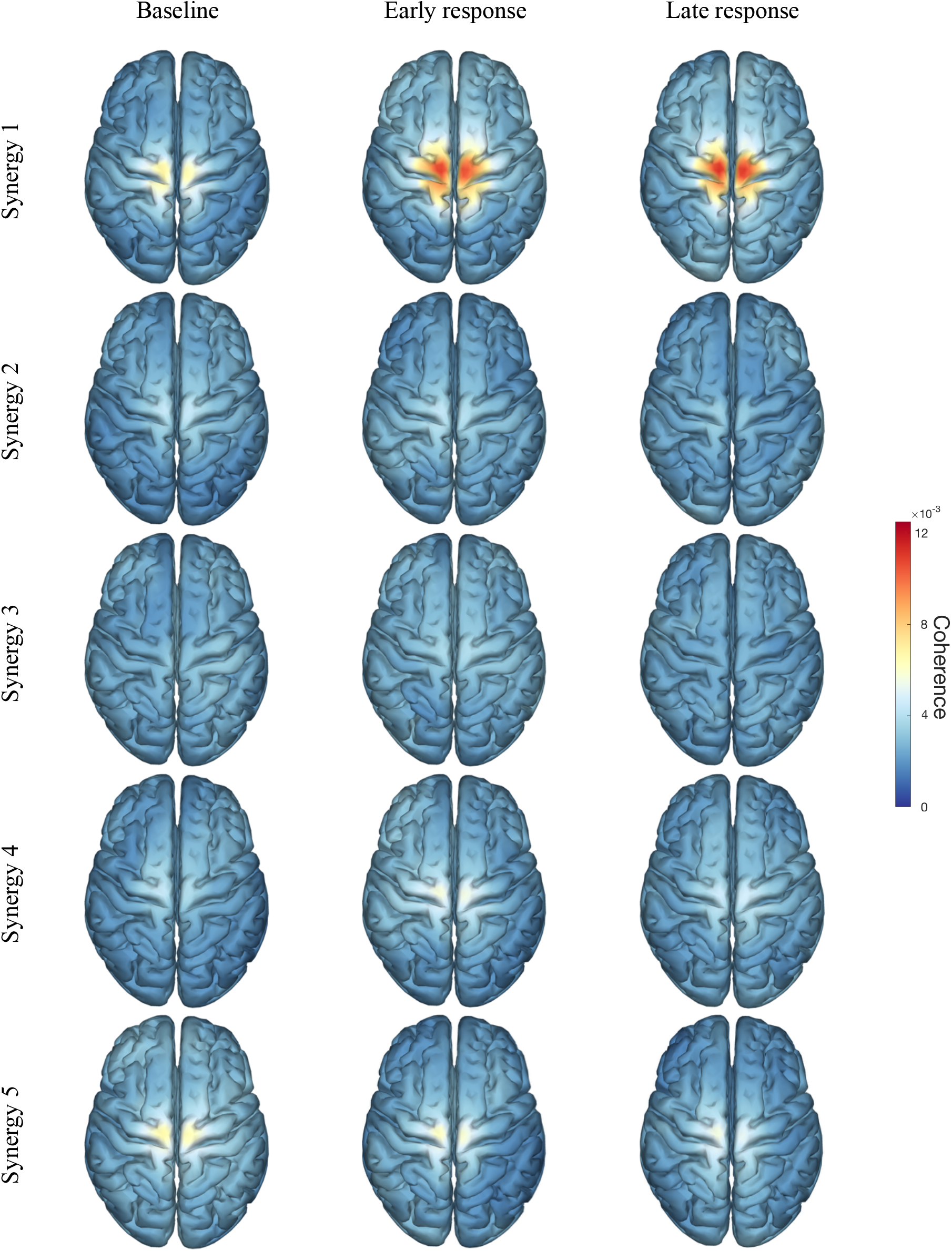
Estimated cortico-synergy coherence during baseline (left column), early response (middle), and late response (right) per synergy. Coherence was pooled over perturbation direction and averaged over sessions. Note that the magnitude of coherence here is higher than in the figures in the body text because they were determined per session to allow for between-session comparison.

### A4 Statistics of cortico-synergy 1 power and coherence per perturbation directions

**Figure A.3.**
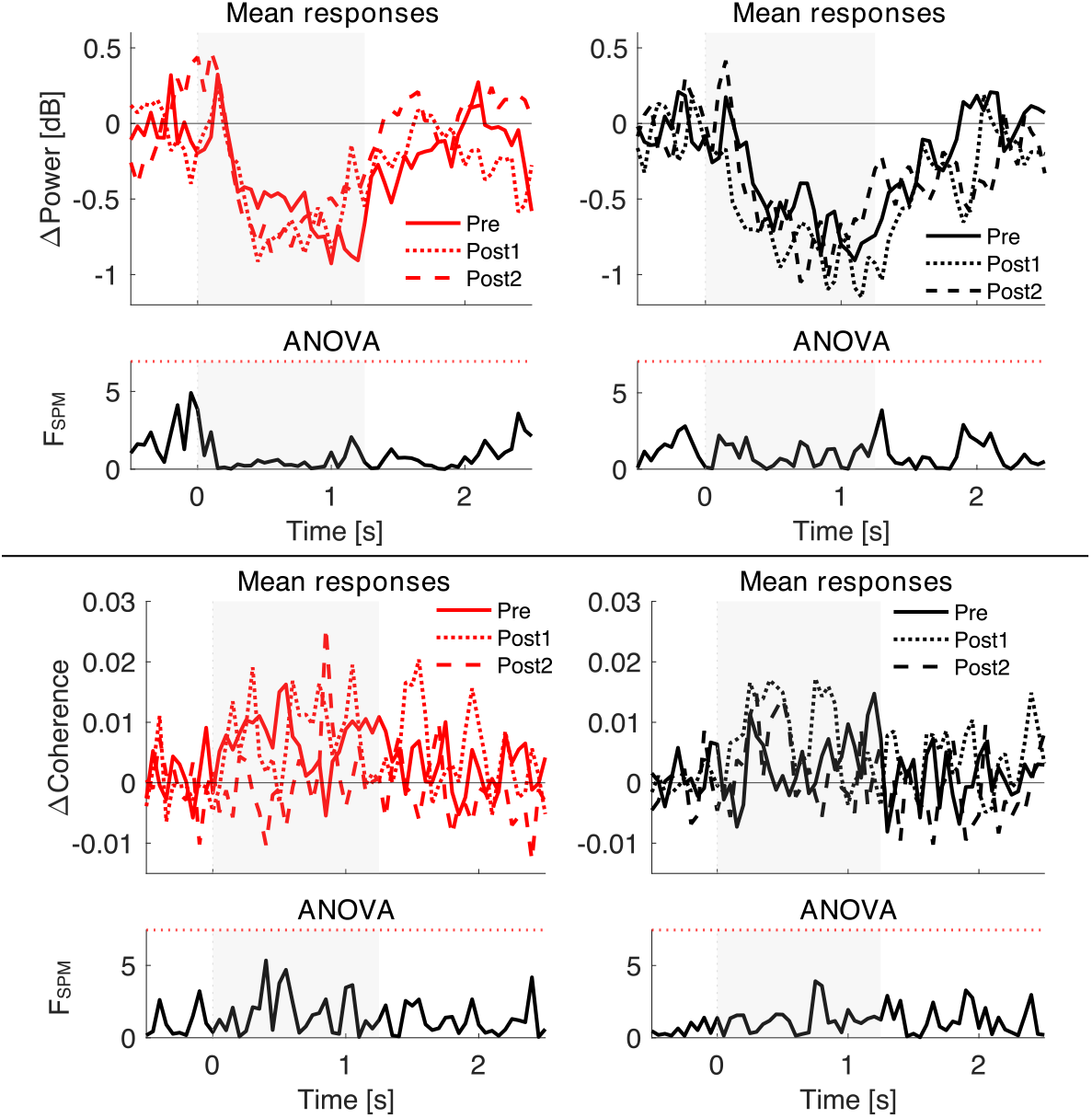
Between sessions comparison of the change in β-power (top panel) and the cortico-synergy coherence (bottom panels). Left column: lateral perturbations; Right column: medial perturbations. Top rows per panel: mean power/coherence per session as a function of time (*t* = 0s ∼ perturbation onset). Bottom rows (per panel): the corresponding *F*-statistics where the red dotted line indicates the corresponding significance threshold. Note that the β-power is expressed as absolute change relative to the baseline period.

## Footnotes

1 Typical candidates for mode decomposition are non-negative matrix factorisation (NMF) or principal or independent component analysis (Tresch et al., 2006). While the first is commonly used because it readily accounts for the semi-positive definiteness of (rectified) muscle activation, it may be biased towards an increased static muscle tonus – an expected effect in our training design. We circumvented this bias by opting for principal components when defining the muscle synergies. Note that the leading principal component, i.e. synergy 1, agrees with the leading independent component rendering differences between PCA and ICA void.

## Notes

### Competing Interest Statement

The authors have declared no competing interest.

